# Proof-of-concept for the detection of early osteoarthritis pathology by endomicroscopy

**DOI:** 10.1101/734368

**Authors:** Mathäus Tschaikowsky, Sofia Brander, Bizan N. Balzer, Bernd Rolauffs, Thorsten Hugel

## Abstract

Osteoarthritis (OA) is a degenerative joint disease and the leading cause of global disability. Clinical trials to date have been unable to pinpoint early and potentially reversible disease states with current clinical technology and hence disease-modifying OA drug candidates cannot be tested early in the disease. To overcome this obstacle, we correlate articular cartilage stiffness changes and cellular spatial organization. The former is a well-understood and functionally relevant OA pathology, while the latter allows discriminating between healthy vs early OA, based on distinct cellular spatial patterns. We demonstrated that an extensive loss of atomic force microscopy-detected stiffness can be seen in cartilage tissues with spatial patterns exhibiting the earliest identifiable OA. In addition, the translation of commercially available clinically usable probe-based confocal laser-endomicroscopy allows us to detect these early OA spatial patterns. This study resolves a major clinical trial obstacle by presenting the proof-of-concept that early OA pathology can be detected by already available clinical technology.

**One Sentence Summary:** We report a correlation between articular cartilage surface nanoscale stiffness and chondrocyte spatial organization; using this correlation enables early pathology detection by currently available clinical optical methods.

## Introduction

*Articular cartilage (AC)* is a specialized tissue that enables joint articulation, lubrication and mechanical loading. It is a biological macromolecular fiber composite material that withstands high tensile, shear and compressive strains while providing very low-friction properties (*1*). The cartilage cells, the chondrocytes, are the tissue’s only resident cell type. They are sparsely distributed across an extra-cellular matrix (ECM) which they maintain. The major structural components of the ECM are a network of collagen fibers and the proteoglycans, which together are the main contributors to AC mechanical properties (*1*). Its overall layered structure consists of four zones, the most superficial zone, constituting the AC surface, the superficial, middle and deep zones (*1–3*) (Fig. 1). Joint lubrication (*4*) and structural adaptations caused by variations in the biomechanical environment primarily occur in the superficial zone (*5*).

**Fig. 1.**
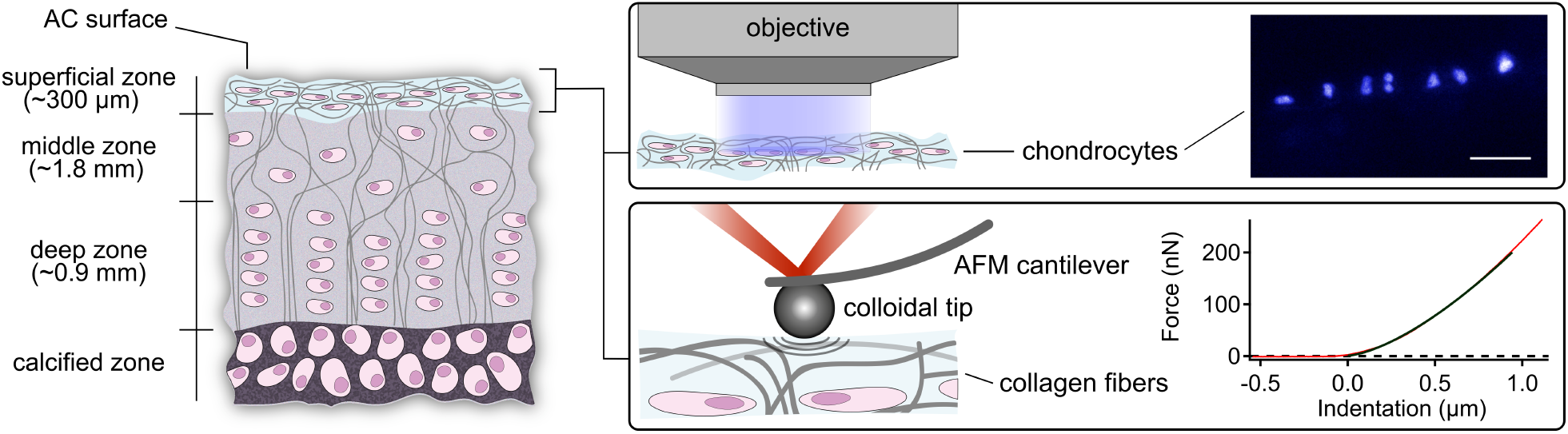
Overview of the sample and the fluorescence and AFM experiments. The left drawing illustrates the various AC layers. The middle drawing zooms into the superficial zone while depicting the utilized methods, namely, fluorescence microscopy of the chondrocytes within the superficial zone (top) and colloidal probe AFM of the AC surface (bottom). The right part of the figure shows at the top a recorded fluorescence microscopy image of human chondrocyte nuclei within the superficial zone that are arranged in a specific pattern of spatial organization termed a double string (top). The chondrocyte nuclei were fluorescent-stained with DAPI and imaged in a top-down view (scale bar corresponds to 50 µm). The bottom right part shows a recorded force vs indentation trace by AFM (red) fitted with the Hertzian contact model (black). Several hundred force vs indentation traces were taken in a grid-like manner (force maps) to obtain an average elastic modulus in three regions of interest for each AC sample. The AFM experiments were carried out with a silicon cantilever and a spherical borosilicate glass particle tip having a radius of 5 (± 0.5) µm. The size of the colloidal particle was chosen to obtain sufficient local resolution while averaging the effects of various structural AC components such as collagen fibers and proteoglycans (see Note S1).

*Osteoarthritis* (OA), the most common joint disorder, is a degenerative disease of the whole joint, for which no preventive or therapeutic biological interventions are available (*6, 7*). OA is characterized by AC degradation and subchondral bone sclerosis, followed by loss of AC and complete destruction of the joint. Research aiming at establishing a unified theory of the initial OA dysfunction has not been successful so far (*6, 8*). The *clinical diagnosis of OA* relies on clinical symptoms and on imaging using x-ray and magnetic resonance imaging (MRI). No diagnostic or prognostic biomarkers are currently able to differentiate healthy vs early OA joints (*9*). MRI scans can acquire morphological and biochemical data (*10*) but have not been able to statistically differentiate between healthy vs early OA (*11*). Other approaches like high frequency ultrasound biomicroscopy, water jet ultrasound indentation and contact mechanical indentation in a controlled animal study (*12*) have not been successful in identifying early OA so far (*13, 14*).

*Atomic force microscopy (AFM)* has been applied to characterize pathological changes of AC several times (*15–21*). It has generally been accepted that AFM is capable of detecting and measuring local structural and functional changes of the AC surfaces (*19*). Therefore, it can locally discriminate between healthy tissue and early OA at the AC surface (*19–21*).

*Fluorescence microscopy* has been used in studies to characterize the spatial organization of superficial zone chondrocytes and for detecting early, preclinical OA (*22–26*). However, so far it has not been known whether the differences in chondrocyte organization within the superficial zone of AC are related to the nanomechanical changes observed in early OA. If so, the next question would be if chondrocyte spatial organization can be detected *in vivo* because this ability would facilitate clinically localizing early OA and estimating the impairment of AC surface nanoscale stiffness.

Fig. 1 shows an overview of this study, which investigates the surface of AC with AFM indentation measurements and the chondrocyte organization of the superficial zone with fluorescence microscopy and finally demonstrates that the chondrocyte organization can be detected *in vivo*. Fresh AC surfaces from human knee joints were horizontally dissected to a final thickness of 300 µm and then mounted in an AFM setup which was positioned on top of a fluorescence microscope platform (see Materials and Methods for details). This enables us to correlate the cantilever tip position, at which AFM indentation data was taken, with the recorded fluorescence microscopy images. AFM indentation experiments allowed us to extract the local stiffness, i.e. elastic moduli, of the samples (*27*).

## Results

### Correlation between AFM-based stiffness and cellular spatial organization, an early OA biomarker

Fig. 2A presents the elastic modulus measured with AFM of human AC from patients with clinical OA, using characterized stages of cellular spatial organization as marker of OA severity. We observe a significant drop in the elastic modulus from more than 200 kPa to less than 50 kPa in AC samples that contain chondrocyte double strings, the earliest identifiable organizational OA marker, compared to AC containing chondrocyte strings, representing AC with healthy cellular spatial organization (Fig. 2B). This indicates that a pronounced loss of AC surface nanoscale stiffness was detectable in those AC samples that were categorized as earliest OA stage, based on their distinct spatial patterns.

**Fig. 2.**
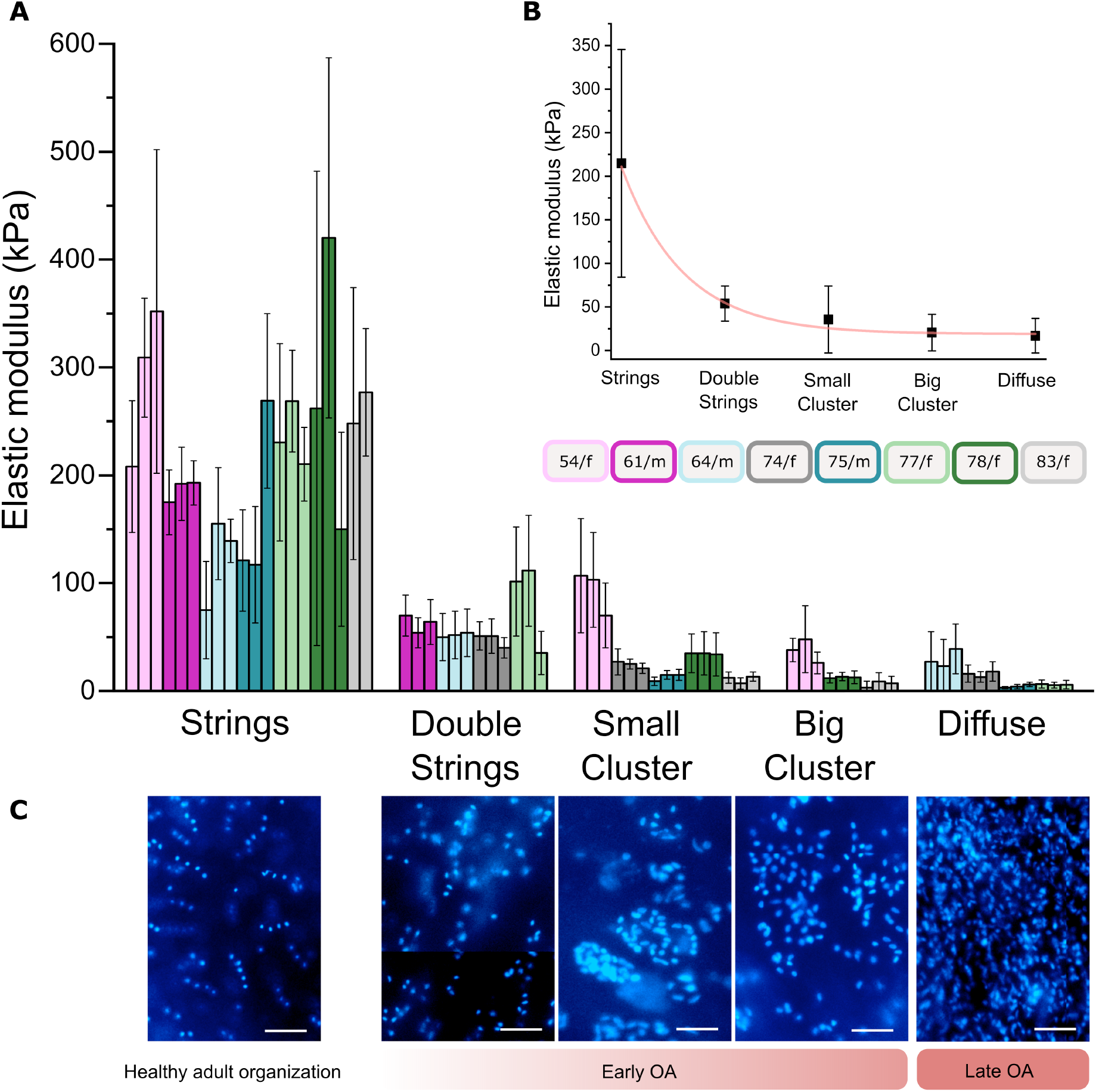
Correlation of fluorescence and AFM-based data. (**A**) Elastic moduli of seven OA patients: Each bar represents the mean elastic modulus of one region of interest (16×16 force vs indentation curves, with standard deviation) corresponding to the classified predominant superficial chondrocyte organization (bars are colour coded with respect to individual patients; boxes indicate age and gender). AFM indentation data show a strong and significant correlation with chondrocyte organization (*r*_*s*_ = −0.68, *p* ≤ 2·10^−7^, number of force vs indentation measurements *n* = 17408). Elastic moduli differences between all types of organization are significant (*p* ≤ 0.001). (**B**), Elastic moduli determined by taking all individual force measurements from all patients corresponding to the classified stage of superficial chondrocyte organization. Number of measurements: strings *n* = 5120; double strings *n* = 3072; small cluster *n* = 3840; big cluster *n* = 2304 and diffuse *n* = 3072. Data were fitted with a single exponential decay. (**C**) Fluorescence images of DAPI stained nuclei of superficial zone chondrocytes displaying distinct stages of healthy (strings) and OA-related spatial organization (double strings to diffuse; the scale bar corresponds to 50 µm).

Prior to AFM measurements the AC samples were fluorescent-stained in order to categorize the samples and use the chondrocyte spatial organization as image-based biomarker. Categorization is based on our lifetime-summarizing model of the cellular spatial organization of a hypothetical human individual (*26*), with the intention to use distinct stages of cellular spatial organization as a marker of OA severity. Fig. 2C presents representative fluorescence images of distinct stages of cellular spatial organization beneath the articular surface, namely strings, double strings, small clusters, clusters, and a diffuse arrangement (see Materials and Methods for sample preparation and classification). Studying the entire AC from a given patient, the AC can contain several stages of cellular spatial organization that vary locally. In this study, we utilized only AC samples that unambiguously displayed one pattern of cellular organization. However, this was not the case for the double string pattern, as strings are known to be interspersed with double strings, indicating an early OA pathology (*24, 26*).

The local elastic moduli of the AC surfaces (superficial zone) significantly correlate with the distinct stages of spatial chondrocyte organization (*r*_*S*_ = −0.68; *p* ≤ 2·10^−7^; number of force vs indentation measurements *n* = 17402; see Note S2 for details). Furthermore, Fig. 2A (and Note S3) indicates that the patient’s age plays a role in the magnitude of the overall decrease of the AC surface elastic modulus: moduli from younger patients remain higher for a given chondrocyte organization.

### Detection of OA stages by clinically applicable confocal laser-endomicroscopy

The above presented results demonstrate that changes in the nanomechanical properties of the AC surface and the OA-induced organizational patterns of superficial zone chondrocytes correlate with each other. Moreover, we are able to pinpoint a major loss of AFM-measured AC surface nanoscale stiffness to the earliest stage of OA, defined here by a specific stage of spatial organization. Based on these data, we envision detection of early OA AC areas with a functionally relevant impairment of AC surface nanoscale stiffness by assessing the cellular spatial organization of the superficial zone, as intra-operative non-destructive AFM measurements do not appear feasible.

In the following we assess probe-based confocal laser-endomicroscopy in this context as a potential tool for improving the currently available methodology to clinically detect early OA. Therefore, we stained chondrocytes of a selected AC explant (Fig. 3A) with calcein and the FDA-approved acriflavine (*28*) *ex vivo* directly after sample preparation on the day of joint-replacement surgery, followed by AC surface imaging (Fig. 3). These experiments demonstrate that we were able to detect different stages of spatial organization of the joint surface chondrocytes, confirming its spatial heterogeneity and the ability of the chosen technology to uncover the tissue’s organizational characteristics for diagnostic purposes. An estimation of local AC surface stiffness is possible using the above presented correlations between nanoscale stiffness and cellular spatial organization (Fig. 3). Movie S1 and Movie S2 present an overview of the several stages that have been visualized within this sample.

**Fig. 3.**
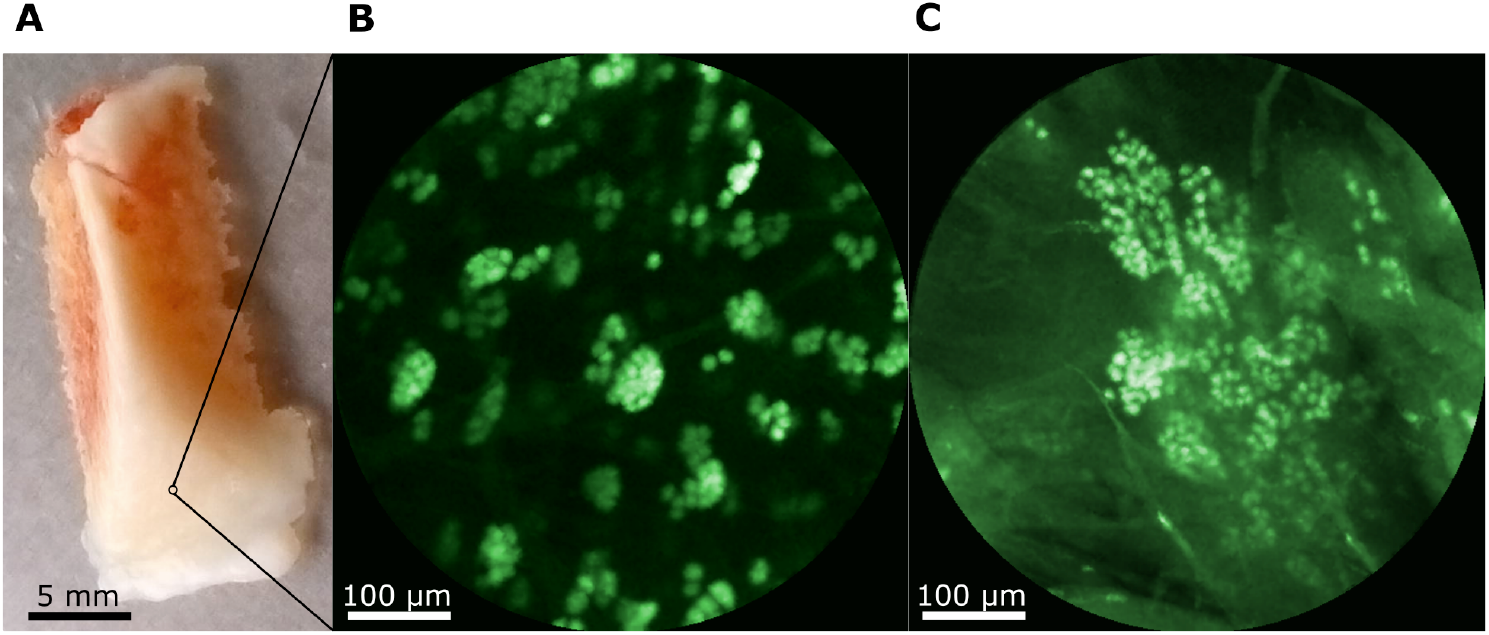
Chondrocyte patterns imaged by ex vivo probe-based confocal laser-endomicroscopy. (**A**) Human AC explant with a region of interest that appears macroscopically intact. (**B**) Human chondrocytes stained with non-toxic calcein (10 µM, 30 min, 37 °C) arranged in small clusters. (**C**) Chondrocytes stained with FDA-approved acriflavine (80 µM, 30 min, 37 °C) displaying a big cluster organization. On the basis of our data and given the age and gender of the patient (female, 68 years), we estimate that the local elastic modulus of the AC surface has decreased by 70 % in (**B**) and 85 % in (**C**) compared to AC with chondrocytes arranged in strings.

## Discussion

By using a combination of fluorescence and AFM-based nanomechanical characterization we demonstrate that the detection of cellular spatial patterns in the superficial zone of AC in early OA correlates significantly with a loss of tissue functionality, namely a reduction of stiffness of the AC surface. This clearly demonstrates that nanomechanical data from human OA AC correlates with a histological score, in particular in early OA. More important, we demonstrate that AC samples categorized by their cellular spatial organization allow for a statistical discrimination between AC samples with intact vs impaired nanomechanical properties. This adds a valuable tool to study the early OA mechanisms because disease onset and progression as well as tissue mechanics can be monitored simultaneously and non-destructively at high resolution with probe-based confocal laser-endomicroscopy. This technique so far has not been used on any musculoskeletal tissues, but is instantly clinically, and more importantly, intra-operatively, applicable.

Endomicroscopy together with the above described correlations between nanoscale mechanics and organizational changes will not only enable clinicians to estimate functional joint surface integrity, but also allows them to monitor AC in animal studies and to diagnose earliest OA in controlled clinical trials, which is currently not possible. Specifically, this point appears valuable, as a series of disappointing late-stage terminations of clinical trials investigating potential disease-modifying OA drugs (DMOADs) has been recorded (*29*) and the design of OA trials has in recent years been a bigger challenge than the identification of novel targets (*30*). Thus, the here introduced methodology adds a powerful tool for early OA diagnostics to the field. This will be particularly helpful for efforts to develop and test DMOADs, because the here introduced approach resolves the insensitivity of the currently available methods to detect early OA.

To discuss the here presented data in a mechanistic context, we speculate that repetitive mechanical trauma to the articular surface, known to lead to microstructural damage, might have contributed to inducing the here observed loss of elastic modulus of AC surface. In addition, this damage might release a proliferation-inducing growth factor such as FGF-2 which is released upon mechanical injury (*31*) to induce changes in the cellular spatial organization of human AC, as demonstrated in (*26*).

Altogether, we provide in this study the necessary data and technical prerequisites to directly enter clinical studies and, therefore, translation of the here presented diagnostic proof-of-concept into the clinic.

## Materials and Methods

### Sample preparation and optical classification

Femoral condyles were obtained from eight patients diagnosed with clinical OA during total knee arthroplasty (ages: 54 - 83 years; 5 female and 3 male patients; see Note S2). All tissues were obtained with approval of the local research ethics committee and informed consent as part of the “Tissue Bank for Research in the Field of Tissue Engineering” project and the biobank “Osteo” (GTE-2002; AN-EK-FRBRG-135/14). The samples were prepared on the day of surgery. Macroscopically intact AC samples were removed from subchondral bone and processed into discs of 2 mm diameter using a biopsy punch (PFM Medical, Germany). The correct orientation of the discs was maintained at all times during further preparation and all experimental procedures. AC surface were horizontally dissected to a final thickness of 300 µm by removing the deeper zones (see Fig. 1). Fluorescent staining was performed with DAPI (1 µg ml^−1^ in PBS; Invitrogen, USA) for 10 min and subsequent washing with PBS. Discs were stored in PBS at 4 °C. Visualization and classification of horizontal chondrocyte patterns were performed 8 to 24 h after explantation. Discs were viewed top-down, i.e. perpendicular to the AC surface. Two-dimensional images capturing the entire cartilage disc were digitally recorded (Axio Observer, Axiocam 506, ZEISS, Germany; magnifications 10x, 20x) and displayed using the ZEN software (ZEISS, Germany). Visual classification was performed according to the sample’s predominant type of chondrocyte organization in the superficial zone by two experienced researchers (*23, 26*). Samples displaying the most unambiguous type of organization were selected for AFM measurements and stored at 4 °C.

### Nanomechanical characterization

The AFM measurements were performed with an MFP-3D AFM (Asylum Research, an Oxford Instruments Company, USA) 48 h after explantation. The AFM head was mounted onto a fluorescence microscope platform (Axio Observer A1, ZEISS, Germany; Clara E camera, Andor, an Oxford Instruments Company, UK). This enabled a correlation of cantilever tip position, force data and local chondrocyte organization in the superficial zone. All experiments were carried out in PBS supplemented with a protease inhibitor cocktail (cOmplete, Roche, Switzerland) at 25 °C. The AC samples were glued onto the glass bottom plate of an AFM fluid cell with a UV curable optical adhesive (OP-4-20632, Dymax, USA) avoiding contamination of the AC surface. A calibration of the AFM cantilever was performed before and after each measurement cycle (see *AFM calibration*). Indentation experiments were carried out with a silicon cantilever with a nominal spring constant of 0.5 to 9.5 N m^−1^ and a spherical borosilicate glass particle having a radius of 5 (± 0.5) µm (CP-FM-BSG-B, sQUBE, Germany). The indentation velocity was 10 µm s^−1^. Three force maps were recorded on each region of interest of the AC sample. Each force map represents a 16 x 16 - grid with a side length of 50 µm and comprising 256 individual points at which force vs indentation curves were taken. These parameters were chosen such that almost the entire area of each force map was probed, given the typical contact area resulting from the radius of the spherical particle and the indentation depth. The samples were measured in a random order to avoid artefacts due to material aggregation on the cantilever tip. The trigger force, i.e. the maximum force that was exerted to indent the sample, was 500 nN. This resulted in a cantilever deflection of approx. 150 nm and a maximum indentation depth of 1 to 6 µm, i.e. 0.3 to 2% of the sample thickness. The indentation depth was chosen to be small compared to the sample thickness (*32, 33*). Force data was analysed in Igor Pro (Wave Metrics, USA) using custom-written software applying the Hertzian contact model (see Note S4). All statistical tests were performed using SigmaStat 12.5 (Systat Software, USA). For details on data analysis see below.

### AFM calibration

Calibration of the AFM cantilever was performed before and after each measurement cycle allowing a conversion of cantilever deflection into force by applying Hooke’s law: first, the inverse optical lever sensitivity (InvOLS) was determined by indenting the cantilever tip into a hard substrate (silicon oxide). Thus, a conversion of the voltage signal from the AFM photodiode into a cantilever deflection could be performed. The spring constant of the cantilever was determined via the thermal noise method (*34, 35*).

### Data analysis and elastic modulus calculation

Force data was analysed in Igor Pro (Wave Metrics, USA) using custom-written software. The elastic modulus *E* for a spherical cantilever tip was computed applying the Hertzian contact model given by eq. 1 (*36, 37*) to the contact region of force extension vs indentation curves:

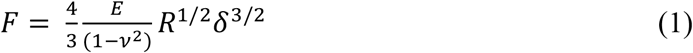

where *F* is the indentation force, ν is the Poisson’s ration of the sample (ν = 0.5 for AC (*38*)), *R* is the radius of the indenter sphere and *δ* is the indentation depth (*δ* ≤ 1 µm).

The Hertzian contact model considers the deformation of an indenting spherical object and a half space with a finite contact area under compression. The model is valid for homogeneous and isotropic materials. Furthermore, it demands the indentation depth to be small compared to the diameter of the ball and no hysteresis side-effects should occur. The maximum indentation depth was less than 2% of the sample thickness, therefore Bueckle’s indentation depth limit of 10% of the sample thickness was not exceeded (*32, 33, 39*). See paragraph "Application of the Hertz model" below for a test and discussion of the other assumptions.

### Statistical analysis

All tests were performed using SigmaStat 12.5 (Systat Software, USA). The errors correspond to the standard deviation of *n* individual measurements, regions of interest (ROIs) or patients. The significance of differences was tested using ANOVA on ranks (*40*) and Dunn’s post-hoc-test(*41*). Additionally, differences between data from samples with string and double string organizations were tested using a two-sample t-test (*42*). Correlation analysis was performed using Spearman’s rank order test (*43*). The resulting correlation coefficient *r*_*S*_ can take values from −1 to 1 corresponding to perfect negative or positive Spearman correlations, respectively. Differences and correlation were considered significant at *p* ≤ 0.05, unless stated otherwise.

### Ex vivo probe-based confocal laser-endomicroscopy

Native and macroscopically intact tissue was obtained during arthroplasty. Chondrocytes were stained *ex vivo* with calcein (10 µM, Invitrogen, USA) and acriflavine (80 µM, Sigma-Aldrich, USA) for 30 min at 37 °C. Samples were imaged with a probe-based confocal laser-endomicroscope (*44*) (Cellvizio, Mauna Kea, France).

### Optical biopsy: Movie S1

This movie shows an optical biopsy of macroscopically intact human AC derived from a knee joint of a joint replacement patient and tissue donor diagnosed with clinical OA (see Fig. 3A).

Chondrocytes were stained *ex vivo* with calcein. Samples were imaged with a probe-based confocal laser-endomicroscope (Cellvizio, Mauna Kea, France). Calcein-stained chondrocytes display different stages of healthy adult and OA-related chondrocyte organization in the superficial zone.

### Optical biopsy: Movie S2

This movie shows an optical biopsy of macroscopically intact human AC derived from a knee joint of a joint replacement patient and tissue donor diagnosed with clinical OA (see Fig. 3A).

Chondrocytes were stained *ex vivo* with acriflavine. Samples were imaged with a probe-based confocal laser-endomicroscope (Cellvizio, Mauna Kea, France). The first part of the video shows acriflavine-stained chondrocytes displaying an OA-related stage of chondrocyte organization in the superficial zone (*big clusters*; 0 to 22 s). Following this, a second region was imaged lacking chondrocytes and thus only displaying the autofluorescence of the AC surface (22 to 27 s). Finally, the probe of the endomicroscope was moved back towards a region showing stained chondrocytes.

## Supplementary Materials

Note S1: Effect of colloidal probe tip size on the elastic modulus.

Note S2: Individual patient data.

Note S3: Effect of age on the decrease of the elastic modulus.

Note S4: Application of the Hertz model.

Movie S1: Optical biopsy of human AC (calcein stain)

Movie S2: Optical biopsy of human AC (acriflavine stain)

## Acknowledgements

This study was in part funded by the Deutsche Forschungsgemeinschaft (DFG, German Research Foundation) under Germany’s Excellence Strategy – EXC-2193/1 – 390951807.

## Author Contributions

B.R. and T.H. designed the study. M.T. performed the experiments with support of S.B. M.T. evaluated the data with support of S.B., B.N.B., B.R. and T.H. The manuscript was written by M.T., B.N.B., B.R., T.H. with support by all authors.

## Data availability

The data that support the findings of this study are available from the corresponding author upon reasonable request.

## References

1. D. W. Smith, B. S. Gardiner, L. Zhang, A. J. Grodzinsky, Articular Cartilage Dynamics (Springer Singapore; Springer, Puchong, Selangor D.E., ed. 1, 2018).

2. E. B. Hunziker, M. Michel, D. Studer, Ultrastructure of adult human articular cartilage matrix after cryotechnical processing. Microsc. Res. Tech. 37, 271–284 (1997).

3. R. Fujioka, T. Aoyama, T. Takakuwa, The layered structure of the articular surface. Osteoarthr. Cartil. 21, 1092–1098 (2013).

4. C. R. Flannery et al., Articular cartilage superficial zone protein (SZP) is homologous to megakaryocyte stimulating factor precursor and Is a multifunctional proteoglycan with potential growth-promoting, cytoprotective, and lubricating properties in cartilage metabolism. Biochem. Biophys. Res. Commun. 254, 535–541 (1999).

5. T. M. Quinn, H.-J. Häuselmann, N. Shintani, E. B. Hunziker, Cell and matrix morphology in articular cartilage from adult human knee and ankle joints suggests depth-associated adaptations to biomechanical and anatomical roles. Osteoarthr. Cartil. 21, 1904–1912 (2013).

6. K. A. Boehme, B. Rolauffs, Onset and Progression of Human Osteoarthritis-Can Growth Factors, Inflammatory Cytokines, or Differential miRNA Expression Concomitantly Induce Proliferation, ECM Degradation, and Inflammation in Articular Cartilage? Int. J. Mol. Sci. 19, 2282 (2018).

7. Q. Liu et al., Knee Symptomatic Osteoarthritis, Walking Disability, NSAIDs Use and All-cause Mortality: Population-based Wuchuan Osteoarthritis Study. Scientific reports. 7 (2017).

8. J. R. Bush, F. Beier, TGF-β and osteoarthritis--the good and the bad. Nat. Med. 19, 667–669 (2013).

9. A. Mobasheri, A.-C. Bay-Jensen, W. E. van Spil, J. Larkin, M. C. Levesque, Osteoarthritis Year in Review 2016: biomarkers (biochemical markers). Osteoarthr. Cartil. 25, 199–208 (2017).

10. L. M. Shapiro et al., Mechanisms of osteoarthritis in the knee: MR imaging appearance. J. Magn. Reson. Imaging. 39, 1346–1356 (2014).

11. B. Bittersohl et al., Zonal T2* and T1Gd assessment of knee joint cartilage in various histological grades of cartilage degeneration: an observational in vitro study. BMJ open. 5, e006895 (2015).

12. Y. Wang, Y.-P. Huang, A. Liu, W. Wan, Y.-P. Zheng, An ultrasound biomicroscopic and water jet ultrasound indentation method for detecting the degenerative changes of articular cartilage in a rabbit model of progressive osteoarthritis. Ultrasound. Med. Biol. 40, 1296–1306 (2014).

13. W. Waldstein et al., OARSI osteoarthritis cartilage histopathology assessment system: A biomechanical evaluation in the human knee. J. Orthop. Res. 34, 135–140 (2016).

14. R. J. H. Custers et al., Reliability, reproducibility and variability of the traditional Histologic/Histochemical Grading System vs the new OARSI Osteoarthritis Cartilage Histopathology Assessment System. Osteoarthr. Cartil. 15, 1241–1248 (2007).

15. S. Kienle et al., Comparison of friction and wear of articular cartilage on different length scales. J. Biomech. 48, 3052–3058 (2015).

16. J. Seog et al., Direct Measurement of Glycosaminoglycan Intermolecular Interactions via High-Resolution Force Spectroscopy. Macromolecules. 35, 5601–5615 (2002).

17. F. P. Rojas et al., Molecular adhesion between cartilage extracellular matrix macromolecules. Biomacromolecules. 15, 772–780 (2014).

18. L. Ng et al., Individual cartilage aggrecan macromolecules and their constituent glycosaminoglycans visualized via atomic force microscopy. J. Struct. Biol. 143, 242–257 (2003).

19. B. Han et al., AFM-Nanomechanical Test. ACS Biomater. Sci. Eng. 3, 2033–2049 (2017).

20. M. Stolz et al., Early detection of aging cartilage and osteoarthritis in mice and patient samples using atomic force microscopy. Nat. Nanotechnol. 4, 186–192 (2009).

21. R. E. Wilusz, S. Zauscher, F. Guilak, Micromechanical mapping of early osteoarthritic changes in the pericellular matrix of human articular cartilage. Osteoarthr. Cartil. 21, 1895–1903 (2013).

22. W. K. Aicher, B. Rolauffs, The spatial organisation of joint surface chondrocytes. Ann. Rheum. Dis. 73, 645–653 (2014).

23. B. Rolauffs, J. M. Williams, A. J. Grodzinsky, K. E. Kuettner, A. A. Cole, Distinct horizontal patterns in the spatial organization of superficial zone chondrocytes of human joints. J. Struct. Biol. 162, 335–344 (2008).

24. B. Rolauffs et al., Proliferative remodeling of the spatial organization of human superficial chondrocytes distant from focal early osteoarthritis. Arthritis. Rheum. 62, 489–498 (2010).

25. B. Rolauffs et al., Onset of preclinical osteoarthritis: the angular spatial organization permits early diagnosis. Arthritis. Rheum. 63, 1637–1647 (2011).

26. T. Felka et al., Loss of spatial organization and destruction of the pericellular matrix in early osteoarthritis in vivo and in a novel in vitro methodology. Osteoarthr. Cartil. 24, 1200–1209 (2016).

27. P. J. Thurner, Atomic force microscopy and indentation force measurement of bone. Wiley Interdiscip Rev. Nanomed. Nanobiotechnol. 1, 624–649 (2009).

28. R. Kiesslich et al., Confocal laser endoscopy for diagnosing intraepithelial neoplasias and colorectal cancer in vivo. Gastroenterology. 127, 706–713 (2004).

29. P. Qvist et al., The disease modifying osteoarthritis drug (DMOAD): Is it in the horizon. Pharmacol. Res. 58, 1–7 (2008).

30. W. Zhang, K. Zou, M. Doherty, Placebos for Knee Osteoarthritis: Reaffirmation of "Needle Is Better Than Pill". Ann. Intern. Med. 163, 392–393 (2015).

31. T. Vincent, M. Hermansson, M. Bolton, R. Wait, J. Saklatvala, Basic FGF mediates an immediate response of articular cartilage to mechanical injury. Proc. Natl. Acad. Sci. U. S. A. 99, 8259–8264 (2002).

32. B. L. Doss, K. Rahmani Eliato, K.-H. Lin, R. Ros, Quantitative mechanical analysis of indentations on layered, soft elastic materials. Soft matter. 15, 1776–1784 (2019).

33. M. Krieg et al., Atomic force microscopy-based mechanobiology. Nat. Rev. Phys. 1, 41–57 (2019).

34. J. L. Hutter, J. Bechhoefer, Calibration of atomic-force microscope tips. Rev. Sci. Instrum. 64, 1868–1873 (1993).

35. J. E. Sader, J. W. M. Chon, P. Mulvaney, Calibration of rectangular atomic force microscope cantilevers. Rev. Sci. Instrum. 70, 3967–3969 (1999).

36. H. R. Hertz, Über die Berührung fester elastischer Körper. J. Reine Angew. Math., 156–171 (1881).

37. F. Rico et al., Probing mechanical properties of living cells by atomic force microscopy with blunted pyramidal cantilever tips. Phys. Rev. E Stat. Nonlin. Soft Matter Phys. 72, 21914 (2005).

38. H. Jin, J. L. Lewis, Determination of Poisson’s Ratio of Articular Cartilage by Indentation Using Different-Sized Indenters. J. Biomech. Eng. 126, 138–145 (2004).

39. H. Bueckle, J. W. Westbrook, H. Conrad (Eds.), The Science of Hardness Testing and Its Research Applications (American Society for Metals, Materials Park, Ohio, 1973).

40. W. H. Kruskal, W. A. Wallis, Use of Ranks in One-Criterion Variance Analysis. J. Am. Stat. Assoc. 47, 583–621 (1952).

41. O. J. Dunn, Multiple Comparisons Using Rank Sums. Technometrics. 6, 241–252 (1964).

42. G. W. Snedecor, W. G. Cochran, Statistical methods (ed. 7, 1982).

43. C. Spearman, The proof and measurement of association between two things. By C. Spearman, 1904. Am. J. Psychol. 100, 441–471 (1987).

44. V. Pavlov et al., Intraoperative Probe-Based Confocal Laser Endomicroscopy in Surgery and Stereotactic Biopsy of Low-Grade and High-Grade Gliomas: A Feasibility Study in Humans. Neurosurgery. 79, 604–612 (2016).

