## Supplementary Materials for "Proof-of-concept for the detection of early osteoarthritis pathology by endomicroscopy"

†shared last authorship.

**Supplementary Materials**

### Note S1:

**Effect of colloidal probe tip size on the elastic modulus**

AFM cantilevers with spherical tips and diameters of 1 µm and 50 nm (biosphere B50-FM/B1000-FM, sQUBE, Germany) were used. The use of a small colloidal probe tip (50 nm) lead to higher average elastic modulus values (see Supplementary Table 1). Furthermore, in a histogram of elastic moduli from diseased AC (*small cluster*) a second population of values appeared (see Supplementary Fig. 1). This can be attributed to force vs indentation curves that were recorded directly on collagen fibres of the exposed network.

**Supplementary Table 1. Effect of different colloidal probe tip sizes on the elastic modulus.** Mean values of elastic moduli as obtained from indentation measurements on healthy (strings) and diseased cartilage (small clusters) with different colloidal probe tip diameters. The error values correspond to the standard deviation. A colloidal probe tip diameter of 10 µm was applied in all other experiments.

| Tip diameter / µm | Strings / kPa | Small Clusters / kPa |
|---|---|---|
| **0.05** | 380 ± 93 | 58 ± 30 |
| **1** | - | 15 ± 8 |
| **10** | 172 ± 57 | 17 ± 9 |

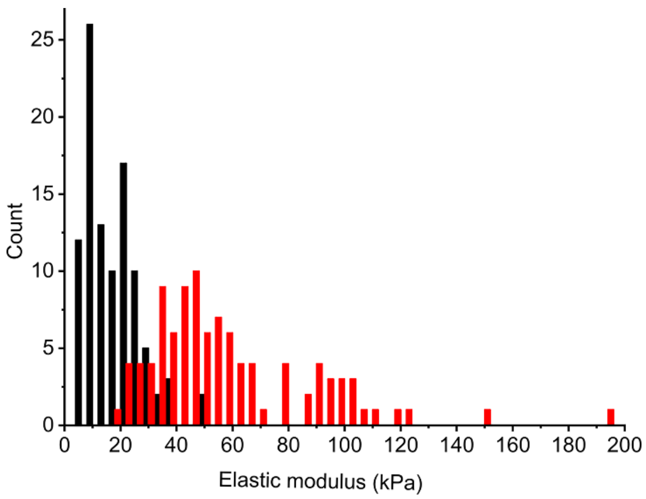


**Supplementary Fig. 1. Effect of different colloidal probe tip sizes on the elastic modulus.** Histograms of elastic moduli calculated from measurements with a small colloidal tip (50 nm diameter; red) and a larger colloidal tip (10 µm diameter; black), respectively. The measurement with the small colloidal tip resulted in a shift of the elastic moduli towards higher values and in an extra population of higher elastic moduli (> 75 kPa).

### Note S2:

#### Individual patient data

Femoral condyles were obtained from eight patients diagnosed with clinical OA during total knee arthroplasty (ages: 54 - 83 years; 5 female and 3 male patients; see Supplementary Table 2 and Supplementary Fig. 2). All tissues were obtained with approval of the local research ethics committee and informed consent as part of the "Tissue Bank for Research in the Field of Tissue Engineering" project and the biobank "Osteo" (GTE-2002; AN-EK-FRBRG- 135/14). The superficial zone was isolated (see Fig. 1). Chondrocytes were stained with DAPI. The chondrocyte arrangement within the native AC was classified in a top-down view according to the predominant type of spatial organization, as published previously (*1*). AFM indentation measurements were performed on three ROIs in a grid-like fashion (16 x 16 force extension vs indentation curves). Elastic moduli were computed using the Hertz model. A significant difference of the elastic modulus between classes of chondrocyte organization can be observed for each individual patient, clearly exceeding the local variation within single samples.

**Supplementary Table 2. Overview for elastic moduli determination.** Total number of patients, ROIs and force vs indentation curves corresponding to each type of chondrocyte organization in the superficial zone. Femoral condyles were obtained from eight patients diagnosed with clinical OA during total knee arthroplasty.

| | Strings | Double strings | Small clusters | Big clusters | Diffuse |
|---|---|---|---|---|---|
| **Patients** | 7 | 4 | 5 | 3 | 4 |
| **ROIs** | 20 | 12 | 15 | 9 | 12 |
| **Force vs indentation curves** | 5120 | 3072 | 3840 | 2304 | 3072 |

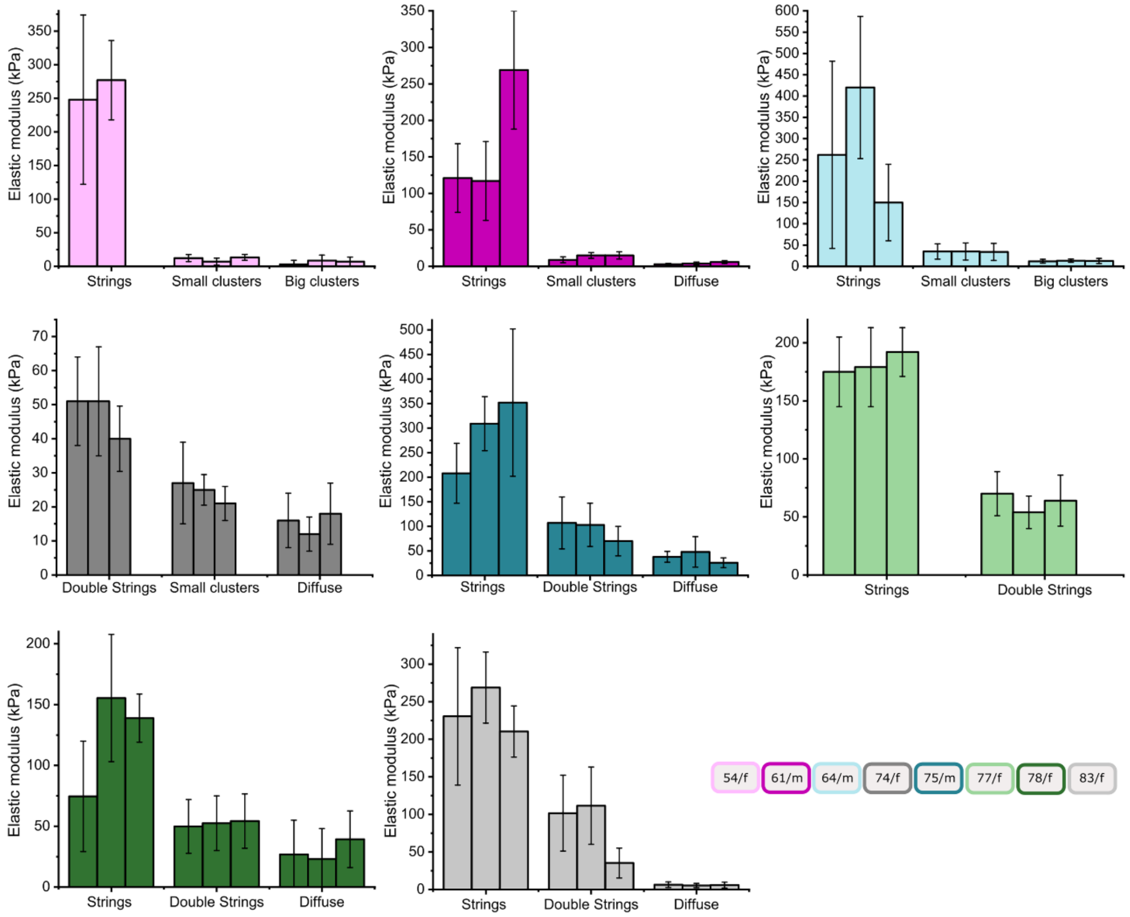


**Supplementary Fig. 2. Correlation of optical and mechanical data.** Each bar represents the mean elastic modulus of one ROI (16 x 16 force vs indentation curves) corresponding to the indicated predominant chondrocyte organization in the superficial zone. The error bars correspond to the standard deviation. Differences in the elastic modulus between all types of organization are significant ($p \leq 0.001$).

**Note S3**

**Effect of age on the decrease of the elastic modulus**

The average elastic moduli from all ROIs corresponding to one type of cell organization were normalized to the value of the respective sample with a predominantly healthy *string*-like organization (see Supplementary Fig. 3; number of patients: $n = 6$). The normalized data reveal a correlation of patient's age and elastic moduli: moduli from patients with highest or lowest elastic modulus in one group remained highest or lowest in another group. The normalized elastic modulus of the surface of diseased AC decreases with age.

Data normalization allowed an insight into a possible age dependence of the decrease of the normalized elastic modulus of the AC surface (see Supplementary Fig. 3). In three out of four groups the oldest patient showed the lowest, while the youngest patient showed the highest normalized elastic modulus.

The normalized elastic moduli show interpatient variations within each group of early-, mid- and late-stage OA-related cell organizations: the interpatient variation was greatest in the late-stage OA cartilage (*diffuse* arrangement).

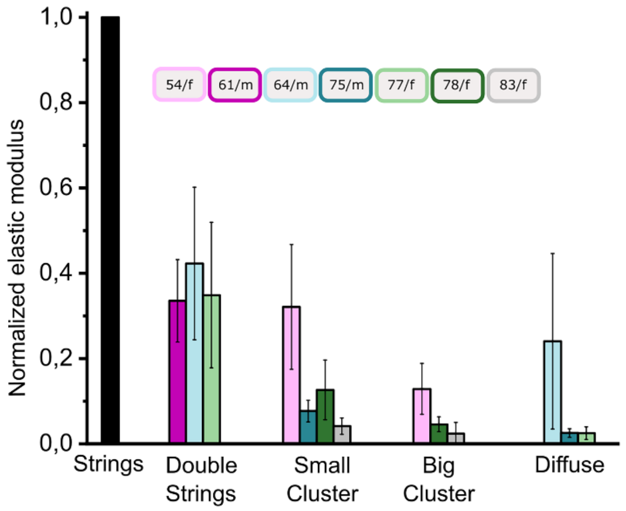


**Supplementary Fig. 3. Effect of age on the elastic modulus.** Elastic moduli for different types of cell organization of all ROIs from each patient are normalized to the value of regions with healthy adult organization of chondrocytes in the superficial zone (strings).

<u>**Note S4**</u>

**Application of the Hertz model**

The application of the Hertz model requires the choice of a contact point, i.e. a starting point for the fit to the force vs indentation curve. The *y*-coordinate of the contact point was set to zero force. The *x*-coordinate (indentation value) of the contact point was determined using a sequential search scheme via $\chi^2$-minimization: starting from a data point near the contact region, a fit for every data point along the *x*-axis was performed. The *x*-coordinate of the data point corresponding to the $\chi^2$-minimum was chosen as the *x*-coordinate of the contact point (*2*).

The applicability of the Hertzian contact model depends on several conditions (*3*). First, the material under investigation must be isotropic and homogeneous. Second, no hysteresis for approach and retraction and no adhesion between cantilever tip and sample should be present. Third, the indentation depth must be small compared to the sphere radius and sample thickness. Regarding material homogeneity, the size of the colloidal probe tip is crucial. AC mainly consists of a network of collagen fibres that is surrounded by a gel-like aggrecan-water layer. Therefore, on a sub-micrometre scale, it is highly inhomogeneous and anisotropic. This was addressed by the choice of an indenter sphere size being large enough to average out the effects of this nanoscale inhomogeneity while preserving a sufficient spatial resolution (see Note S1).

Several experimental approaches were performed to support the use of the Hertz model:

a. indentation experiments testing for the plasticity of the AC surface, where force vs indentation curves were recorded at a constant *x*-*y*-position and the change of the contact point position was examined.

b. application of cantilever tip velocities of 0.5 - 20 $\mu m\ s^{-1}$ testing for the rate-dependence of the elastic modulus determined by nanoindentation of human AC.

c. application of the DMT model to a AC surface dataset for comparison to the Hertz model using a commercially available software (Asylum Research, an Oxford Instruments Company, USA) (*4*).

**Indentation experiments testing for the plasticity of the AC surface**

A weak hysteresis and adhesion were observed in most force vs indentation curves (see Supplementary Fig. 4a), but the error due to this hysteresis is smaller than the error resulting from AC surface inhomogeneity and is therefore negligible. In order to test for plasticity repeated force vs indentation curves were recorded at a constant *x*-*y*-position monitoring the change of the contact point position (see Supplementary Fig. 4b). A small drift of the contact point was observed. Three out of four experiments indicate a plastic deformation of the AC surface of approximately 1.5 nm per force vs indentation curve in *z*-direction. This is very little compared to the micrometre indentation and is therefore negligible.

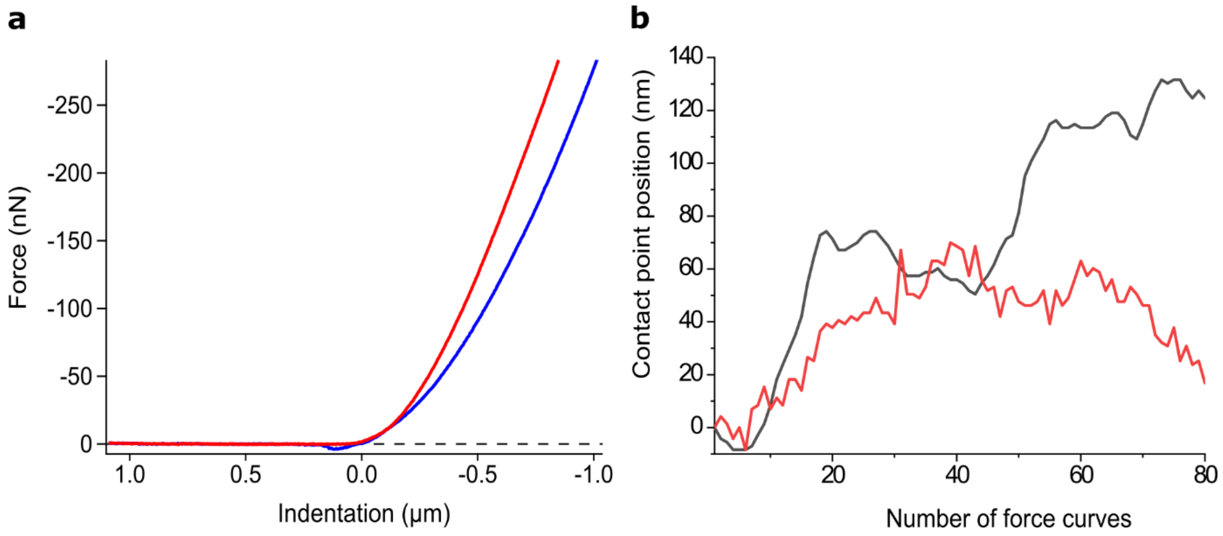


**Supplementary Fig. 4. Test for plasticity.** (**a**) Force vs indentation curve: approach (red) and retraction part (blue) showing an adhesion peak height of 3.7 nN in the retraction part and a slight hysteresis. (**b**) Contact point position with respect to the initial contact point for repeated indentation at a constant x-y-position: Two representative measurements (black and red curves) on different AC samples taken from one patient.

#### a) Rate-dependence of the elastic modulus of the AC surface

The rate-dependence of the elastic modulus was probed. Different cantilever tip velocities were applied (see Supplementary Table 3). A 20-fold decrease in velocity resulted in a 13% decrease of the average elastic modulus. A doubling of the velocity increased the average elastic modulus by 5%. This indicated a slight rate-dependence of the elastic modulus determined by nanoindentation of the AC surface. In order to further minimize rate-dependent effects all experiments presented in the main text have been taken with 10 µm $s^{-1}$.

**Supplementary Table 3. Rate-dependence of the elastic modulus.** The Mean elastic moduli obtained from measurements with varying cantilever tip velocities show a slight rate-dependence. The error values correspond to the standard deviation.

| **Indentation velocity / µm $s^{-1}$** | **Mean elastic modulus / kPa** |
|---|---|
| **0.5** | 121 ± 22 |
| **10** | 138 ± 20 |
| **20** | 146 ± 22 |

### b) Application of DMT model

Finally, to further validate the use of the Hertz model for human osteoarthritic cartilage, a sample dataset was fitted using the DMT model (*4*). The DMT model assumes the same contact profile as the Hertzian approach. In addition to the Hertz model it considers an adhesion force that results from adhesion between the cantilever tip and an underlying surface which results in an adhesion peak in the retraction part of the force vs indentation curve. Another common contact model is the JKR model (*5*), which is only valid for strong adhesion and therefore was not applied.

The force values of this adhesion peak for the obtained data (see Supplementary Fig. 4a) are negligible because their consideration led to elastic moduli values that were similar to values obtained from applying the Hertz model (see Supplementary Table 4). The Oliver-Pharr method (*6*) was also tested and qualitatively similar results were obtained. Thus, the Hertz model was applied for all other experiments.

**Supplementary Table 4. Elastic moduli obtained by application of different contact models.** Mean elastic moduli as obtained by the Hertz and the DMT model. The error values correspond to the standard deviation.

| Contact model | Strings / kPa | Small Clusters / kPa |
|---|---|---|
| **Hertz** | 138 ± 38 | 43 ± 4 |
| **DMT** | 127 ± 26 | 44 ± 8 |

**Supplementary References**


1. T. Felka *et al.,* Loss of spatial organization and destruction of the pericellular matrix in early osteoarthritis in vivo and in a novel in vitro methodology. *Osteoarthr. Cartil.* **24**, 1200–1209 (2016).

2. D. C. Lin, E. K. Dimitriadis, F. Horkay, Robust strategies for automated AFM force curve analysis--I. Non-adhesive indentation of soft, inhomogeneous materials. *J. Biomech. Eng.* **129**, 430–440 (2007).

3. M. Scherge, S. N. Gorb, *Biological micro- and nanotribology*, *Nature's solutions* (Springer, Berlin, 2001).

4. B.V. Derjaguin, Muller, V.M, Toporov, Yu.P, Effect of contact deformations on the adhesion of particles. *J. Colloid Interface Sci.* **53**, 314–326 (1975).

5. K. L. Johnson, K. Kendall, A. D. Roberts, Surface Energy and the Contact of Elastic Solids. *Proc. Royal Soc. A*. **324**, 301–313 (1971).

6. W. C. Oliver, G. M. Pharr, An improved technique for determining hardness and elastic modulus using load and displacement sensing indentation experiments. *J. Mater. Res.* **7**, 1564–1583 (1992).
